# UXS1 regulates UDP-GlcA levels to support growth of UGDH-high cancer cells

**DOI:** 10.1101/2024.02.08.579288

**Authors:** Yunyi Wang, Chen Zhou, Berna Bou-Tayeh, Richard Lisle, Skirmantas Kriaucionis, Yang Shi

## Abstract

Identifying genes that are crucial for cancer cell survival but dispensable in normal cells holds immense therapeutic potential. The DepMap Consortium’s extensive datasets have paved the way for uncovering such selectively essential genes in cancer. However, it remained challenging to efficiently prioritize understudied, selectively essential genes for validation and characterization. To this end, our lab has previously ranked and prioritized potentially understudied, selectively essential genes based on their PubMed publication numbers. This approach led to successful identification and detailed characterization of two top understudied genes. Building on this methodology, our current research identified UXS1, an enzyme responsible for catalyzing the conversion from UDP-glucuronic acid (UDP-GlcA) to UDP-xylose, as a selectively essential gene in cancer cells expressing elevated levels of UGDH, an enzyme responsible for producing UDP-GlcA. Through an integrated approach combining genetic and biochemical assays, we discovered that UXS1 plays a critical role in these UGDH-overexpressing cancer cells by preventing the harmful buildup of UDP-GlcA, which otherwise would lead to cellular toxicity. Our findings not only validate our strategy for prioritizing underexplored but potentially pivotal selectively essential genes but also highlight UXS1 as a potential vulnerability and therapeutic target in cancers characterized by high UGDH expression.

---

Genes selectively essential in cancer cells but not normal cells are optimal cancer therapeutic targets. Due to limited gene essentiality information in normal cells, exploring genes selectively essential in a subset of cancers, but not across all cancers, serves as a practical starting point for uncovering potential therapeutic targets. These selectively essential genes can be identified from the DepMap portal, where systematic genome-wide CRISPR/Cas9-based fitness screening experiments were performed with more than 1000 cancer cell lines originated from a wide range of cancer types (Tsherniak et al., 2017). In fact, many therapeutic targets of FDA-approved cancer drugs are defined as selectively essential in the DepMap portal while many cancer drugs that failed in clinical trials target common essential genes (Chang et al., 2021), suggesting bona fide cancer therapeutic targets may be identified from selectively essential genes. Previously, our lab collected 347 selectively essential genes from the DepMap portal based on the NormLRT score method (McDonald et al., 2017) and prioritized understudied ones by using the number of PubMed publications. This allowed us to identify two unknown genes, C11orf53 (POU2AF2) and COLCA2 (POU2AF3), as selective vulnerabilities in POU2F3-dependent tuft cell-like small cell lung cancer and demonstrated, as did two other publications (Szczepanski et al., 2022; Wu et al., 2022), that they function as transcription co-activators for POU2F3 (Zhou et al., 2022), demonstrating the effectiveness of our strategy to identify novel cancer vulnerabilities.

In addition to C11orf53 and COLCA2, we identified UXS1 (UDP-glucuronate decarboxylase 1) as one of the most understudied in our collection of selectively essential genes. We prioritized UXS1 for two key reasons. First, the distribution of UXS1 CERES scores, a measurement of the effect of gene knockout on cell growth where zero indicates no effect and -1 is the median for common essential genes, across more than 1000 cancer cell lines demonstrated its selective essentiality in only a subset of cancer cell lines (Fig. 1a). In addition, the CERES scores of UXS1 in some cell lines were less than -1, suggesting strong growth defects upon loss of UXS1 (Fig. 1a). Second, one of the top associations pre-computed by the DepMap portal was a negative correlation between the CERES scores of UXS1 and UGDH (UDP-glucose 6-dehydrogenase) expression levels (Fig. 1b, Supplementary Fig. S1). This would imply that UXS1 is selectively essential in cancer cells expressing high levels of UGDH, which is in the same biochemical pathway as UXS1 (Fig. 1c). UGDH is the sole enzyme in human that converts uridine diphosphate (UDP)-glucose to UDP-glucuronic acid (UDP-GlcA), which can be converted to UDP-xylose by UXS1. UDP-xylose serves mainly as a substrate in the synthesis of proteoglycans, which are involved in a number of signaling pathways especially during development. On the other hand, UDP-GlcA participates in three major downstream pathways: synthesis of proteoglycans, production of hyaluronan, as well as glucuronidation (Moriarity et al., 2002; Zimmer et al., 2021). While the involvement of UXS1 in cancer was explored, its impact remained incompletely understood. One study showed that UXS1 knockdown in a prostate cancer cell line resulted in only a transient increase in cell growth rate (Zimmer et al., 2016) and another study reported that depletion of UXS1 in breast cancer cells had no significant effect on tumor growth or lung metastasis (Teoh et al., 2020). Consequently, the significance of UXS1 in cancer remained unclear.

**Fig. 1.**
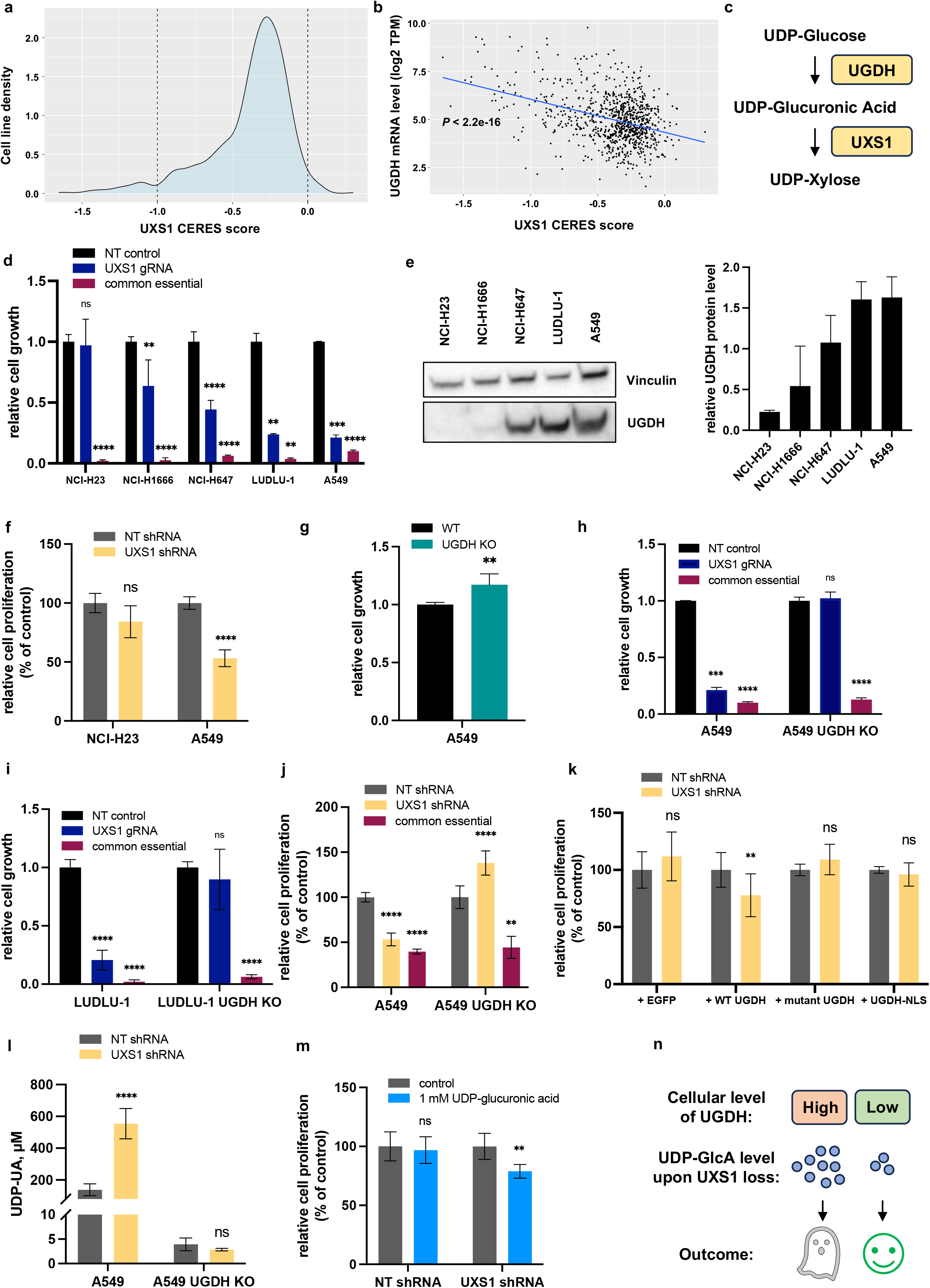
UXS1 regulates UDP-GlcA levels to support growth of UGDH-high cancer cells. **a**. Density plot representing the distribution of UXS1 CERES scores in cancer cell lines. DepMap Public 21Q2 dataset was used. **b**. Scatter plot showing the relationship between UGDH expression level and UXS1 dependency in cancer cell lines. DepMap Public 21Q2 dataset was used. **c**. Diagram showing the biochemical pathway involving UGDH and UXS1. **d**. Relative cell growth, measured with CyQUANT cell proliferation assay on day 0 and day 8, of cancer cell lines treated with NT (non-targeting) gRNA, UXS1 gRNA or PSMB3 (common essential) gRNA. **e**. UGDH protein levels in different cancer cell lines. Western blotting data were quantified. **f**. Relative cell proliferation, measured with BrdU assay, of NCI-H23 and A549 cells treated with NT shRNA or UXS1 shRNA. **g**. Relative cell growth, measured with CyQUANT cell proliferation assay, of WT (wildtype) and UGDH KO A549 cells. **h**. Relative cell growth, measured with CyQUANT cell proliferation assay on day 0 and day 8, of WT and UGDH KO A549 cells treated with NT gRNA, UXS1 gRNA or PSMB3 (common essential) gRNA. **i**. Relative cell growth, measured with CyQUANT cell proliferation assay on day 0 and day 8, of WT and UGDH KO LUDLU-1 cells treated with NT gRNA, UXS1 gRNA or PSMB3 (common essential) gRNA. **j**. Relative cell proliferation, measured with BrdU assay, of WT and UGDH KO A549 cells treated with NT shRNA, UXS1 shRNA or PSMB3 (common essential) shRNA. **k**. Relative cell proliferation, measured with BrdU assay, of UGDH KO A549 cells overexpressing EGFP/WT UGDH/mutant UGDH/UGDH-NLS (nuclear localization signal) and treated with NT or UXS1 shRNA. **l**. Intracellular UDP-GlcA level measured with HPLC-MS/MS in WT/UGDH KO A549 cells treated with NT/UXS1 shRNA. **m**. Relative cell proliferation, measured with BrdU assay, of UGDH KO A549 cells expressing NT/UXS1 shRNA and treated with/without exogenous UDP-GlcA. **n**. Diagram showing the UDP-GlcA toxicity model. Data are mean ± s.d. Two-tailed unpaired Welch’s t-test was performed for all statistical analysis. ns, not significant. *P<0.05; **P<0.01;***P < 0.001; ****P < 0.0001.

First, we set out to confirm the selective UXS1 dependency in cancer cell lines showing high UGDH expression. We used the CRISPR/Cas9 system to knock out UXS1 in five cancer cell lines with different UXS1 dependency and UGDH mRNA levels as indicated by the DepMap data (Supplementary Fig. S2). The growth suppression effect of UXS1 deficiency was correlated with UGDH protein expression levels, with higher growth suppression seen in cell lines with higher UGDH protein levels (Fig. 1d, 1e). The differential growth suppression effect was not due to differential CRISPR/Cas9 knockout efficiency (Supplementary Fig. S3). We also performed shRNA knockdown experiments and confirmed the stronger requirement for UXS1 in A549 cells, exhibiting the highest UGDH level among the five cell lines tested, than in NCI-H23, which had the lowest UGDH level (Fig. 1f, Supplementary Fig. S4). In addition, exogenous expression of shRNA-resistant UXS1 successfully rescued the growth defects of A549 cells (Supplementary Fig. S5), ruling out potential off-target effects of the UXS1 shRNA. Therefore, consistent with the data in the DepMap portal, UXS1 is indeed selectively essential in cancer cell lines with high UGDH levels.

In order to explain the UXS1 dependency in cell lines expressing high levels of UGDH, we proposed two hypotheses. In the first one, cell lines expressing high levels of UGDH might require more UDP-xylose or its downstream derivatives for growth. According to the first hypothesis, loss of UGDH, which can reduce UDP-xylose levels in the same way as loss of UXS1, should be essential in UXS1-dependent cell lines. However, UXS1-dependent cancer cells did not have very negative UGDH CERES scores in the DepMap portal (Supplementary Fig. S6), indicating that UGDH might be dispensable in the cells dependent on UXS1. In line with this, knockout of UGDH in A549, a UXS1-dependent cell line, did not suppress its growth (Fig. 1g, Supplementary Fig. S7). Thus, lack of UDP-xylose or downstream products could not account for the requirement of UXS1 in UGDH-high cells.

Next, we hypothesized that loss of UXS1 might cause overaccumulation of the upstream metabolite, UDP-GlcA, only in cells with high levels of UGDH, which can cause cell toxicity. To test this hypothesis, we examined UGDH knockout (KO) A549 cells’ sensitivity to loss of UXS1. We showed that depletion of UGDH protected the cells from the growth suppression caused by UXS1 KO (Fig. 1h, Supplementary Fig. S7, S8). Similar results were obtained in UGDH KO LUDLU-1 cells (which also highly express UGDH) treated with UXS1 gRNA (Fig. 1i, Supplementary Fig. S7, S8) and UGDH KO A549 cells treated with UXS1 shRNA (Fig. 1j, Supplementary Fig. S4, S7). To confirm that the lack of UDP-GlcA overaccumulation rather than the loss of other potential functions of UGDH in the UGDH KO cells accounted for the rescue phenotype, we reintroduced wildtype or catalytic-dead UGDH into UGDH KO cells and evaluated their dependency on UXS1. As expected, re-expression of wildtype but not catalytic-mutant UGDH re-sensitized the UGDH KO A549 cells to UXS1 loss (Fig. 1k, Supplementary Fig. S9). Since we and others (Wang et al., 2019) found that UGDH is localized in both nucleus and cytoplasm (Supplementary Fig. S10) in wildtype A549, we also introduced nucleus-targeting UGDH into UGDH KO A549 cells. We found that the exclusively nucleus-localized UGDH did not restore sensitivity of UGDH KO A549 cells to UXS1 loss, suggesting that the catalytic function of UGDH in the cytoplasm is likely to be responsible for the phenotype. We also measured the intracellular levels of UDP-GlcA in wildtype and UGDH KO A549 cells with or without UXS1 knockdown. Consistent with the hypothesis, intracellular UDP-GlcA levels increased dramatically upon UXS1 knockdown in wildtype but not in UGDH KO cells (Fig. 1l). In accordance with this observation, adding 1 mM UDP-GlcA directly to UGDH KO A549 cells reduced cell proliferation only when UXS1 is knocked down (Fig. 1m). This further confirms that the overaccumulation of the metabolite is toxic to the cells and UGDH-high cancer cells depend on UXS1 to prevent excessive accumulation of UDP-GlcA.

In summary, we leveraged the data in the DepMap portal and validated UXS1 as a selectively essential gene in cancer cell lines expressing high levels of UGDH. We also demonstrated that in UGDH-high cancer cells, excessive accumulation of UDP-GlcA after loss of UXS1 was toxic and likely to be the reason for the selective essentiality of UXS1 in UGDH-high cancer cells (Fig. 1n). This conclusion was also reached by a comprehensive, in-depth study recently published in Nature (Doshi et al., 2023). As discussed earlier, our study additionally investigated the role of the nucleus localized UGDH in this process, and found that nuclear UGDH did not contribute to UXS1 sensitivity. However, it is important to note that nuclear UGDH may play a role in other facets of cancer development and progression (Hagiuda et al., 2019). Collectively, these findings suggest that UXS1 might be a promising therapeutic target for cancers showing high UGDH expression and UGDH might be a valuable biomarker. This has broad implications for cancer treatment as high UGDH expression is not limited to specific types of cancer (Supplementary Fig. S11) (Li et al., 2020) and high UGDH expression is associated with worse overall survival in cancer patients (Supplementary Fig. S12) (Tang et al., 2019). However, considering that UXS1^-/-^ mice are not viable (Tang et al., 2010) and that some normal cells also express high levels of UGDH (Supplementary Fig. S11), potential side effects of targeting UXS1 should be monitored carefully. Of note, our discovery of UDP-GlcA overaccumulation as a cause of cellular toxicity aligns with the “kitchen sink” model that has been proposed in cancer metabolism (Lee et al., 2020). According to this model, when an upstream enzyme excessively produces a metabolite, akin to a water tap turned on excessively, the downstream enzyme functions like a drain to reduce the levels of the potentially toxic metabolites. Consistent with this model, SEPHS2 has been reported to be essential in selenophilic cancers to process toxic selenide (Carlisle et al., 2020). Therefore, we speculate that more metabolic vulnerabilities of cancer that can prevent overaccumulation of toxic metabolites will be unveiled in the future.

## Acknowledgements

We acknowledge the Broad Institute for making the data on DepMap portal available to the research community. This project is supported by the Ludwig Institute for Cancer Research. Y.S. is an American Cancer Society Research Professor.

## Author contributions

C.Z. and Y.S. conceived the initial project and wrote the manuscript with input from all the co-authors. Y.W. designed and performed most of the experiments. C.Z. performed bioinformatic analysis. B.B.T helped with cloning and CyQUANT assays, and participated in the discussions during the course of the study. S.K. performed metabolites measurement experiments. R.L. helped with immunostaining experiments.

## Conflict of interest

Y.S. is a co-founder of K36 Therapeutics and ABio Inc, and a member of ABio’s Board of Directors. Y.S. is also a member of the Scientific Advisory Board of EPICRISPR BIOTECHNOLOGIES, INC. Y.S. holds equity in Active Motif, K36 Therapeutics and ABio, Inc. All other authors declare no competing interests.

## Notes

### Competing Interest Statement

Y.S. is a co-founder of K36 Therapeutics and ABio Inc, and a member of ABio Board of Directors. Y.S. is also a member of the Scientific Advisory Board of EPICRISPR BIOTECHNOLOGIES, INC. Y.S. holds equity in Active Motif, K36 Therapeutics and ABio, Inc. All other authors declare no competing interests.

