## Supplementary Materials for "UXS1 regulates UDP-GlcA levels to support growth of UGDH-high cancer cells"

### Supplementary Fig. S1

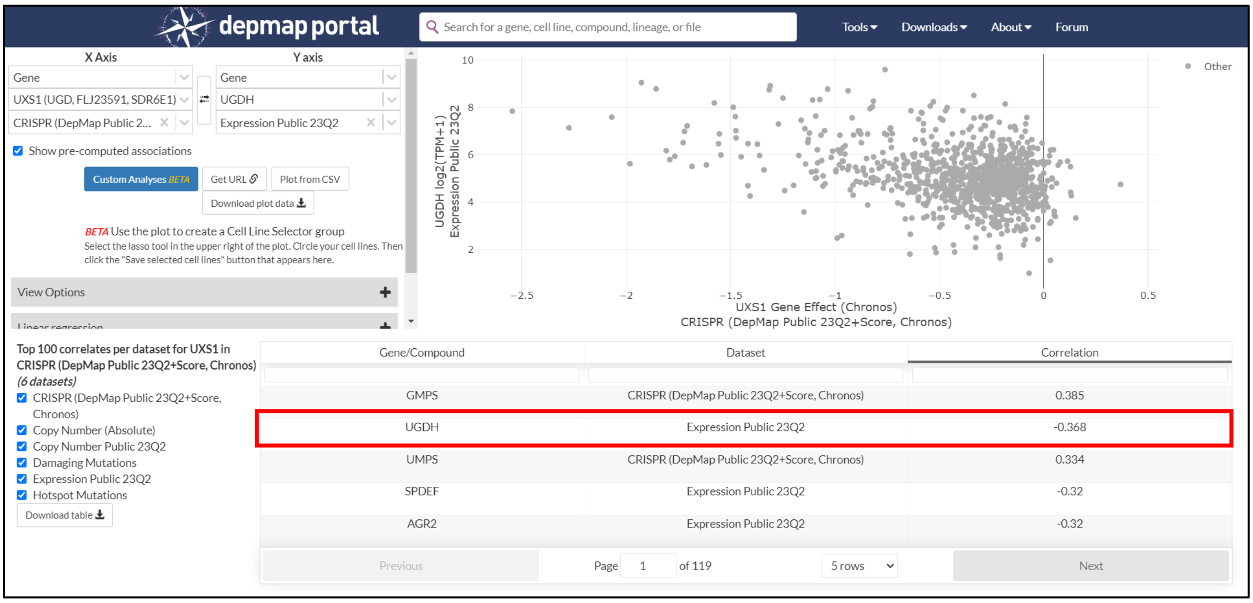


**Supplementary Fig. S1 UGDH expression is the second top pre-computed correlate for UXS1 dependency in the DepMap portal.** Screenshot of the data explorer tab in the DepMap portal. Top 100 correlates for UXS1 dependency were pre-computed and provided by the DepMap portal.

### Supplementary Fig. S2

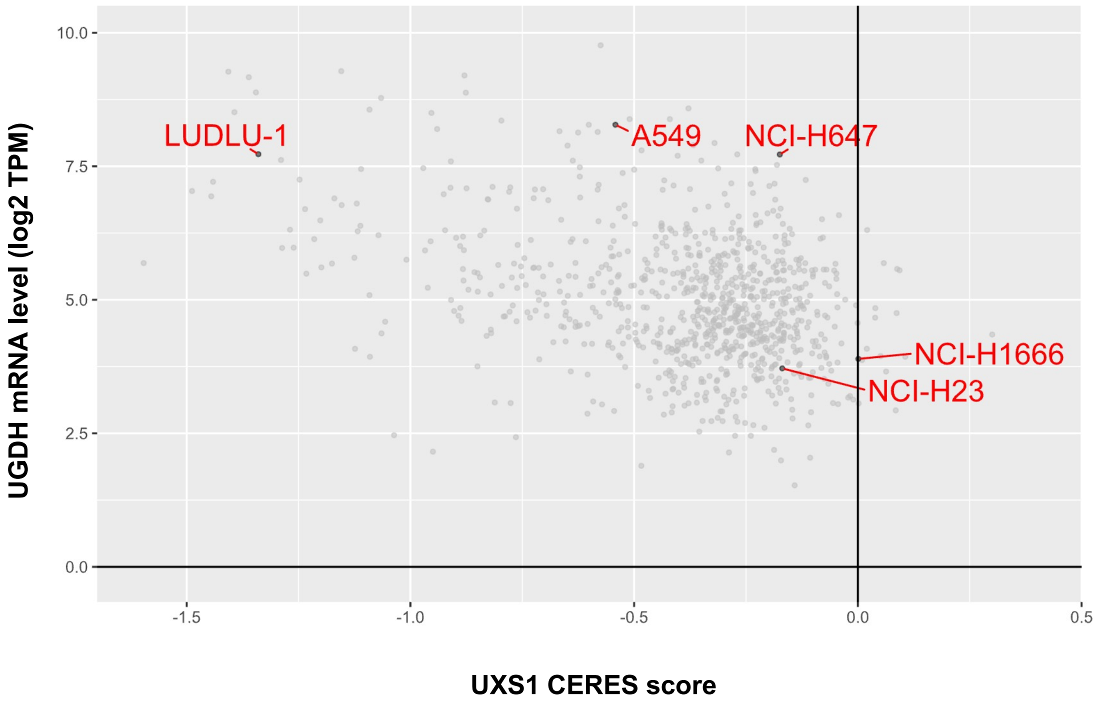


**Supplementary Fig. S2 Cancer cell lines selected for UXS1 dependency testing.** Scatter plot showing the relationship between UGDH mRNA level and UXS1 CERES score in cancer cell lines. Five cancer cell lines were selected for validation. Figure was generated with DepMap Public 21Q2 data.

**Supplementary Fig. S3**

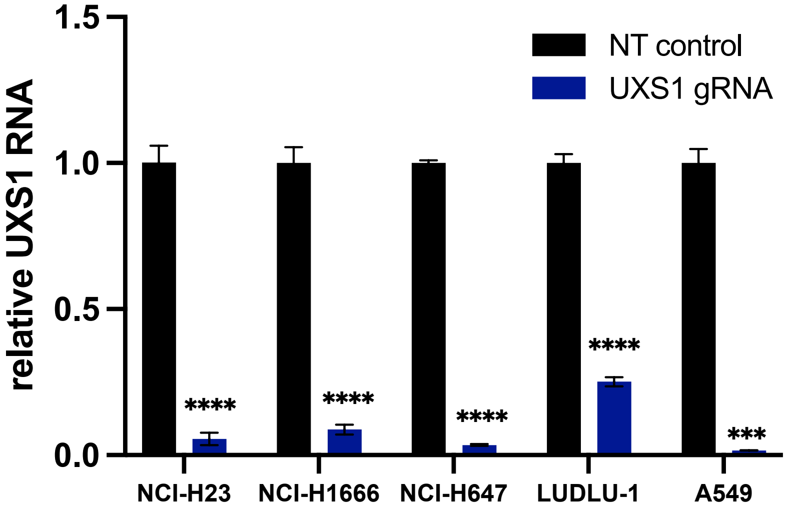


**Supplementary Fig. S3 Similar CRISPR KO efficiency in different cancer cell lines.** UXS1 mRNA levels were measured by RT-qPCR in wildtype and UXS1 KO cancer cell lines. Data are mean ± s.d. Two-tailed unpaired Welch's t-test was performed. ***P < 0.001; ****P < 0.0001.

**Supplementary Fig. S4**

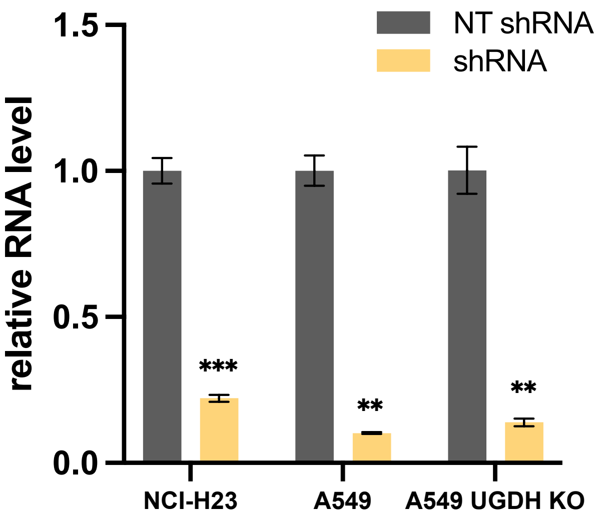


**Supplementary Fig. S4 UXS1 was successfully knocked down with shRNA.** UXS1 mRNA levels were measured by RT-qPCR in wildtype NCI-H23, wildtype A549 and UGDH KO A549. Data are mean ± s.d. Two-tailed unpaired Welch's t-test was performed. $**P < 0.01$; $***P < 0.001$.

**Supplementary Fig. S5**

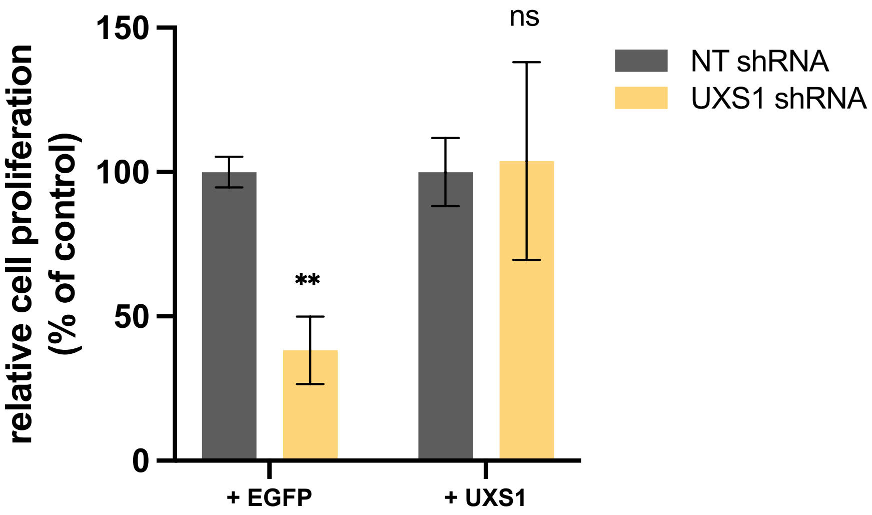


**Supplementary Fig. S5 shRNA-resistant UXS1 rescued A549 cells from growth defects due to UXS1 loss.** Bar plot showing relative growth of A549 cells expressing EGFP or shRNA-resistant UXS1 and treated with NT/UXS1 shRNA. Data are mean ± s.d. Two-tailed unpaired Welch's t-test was performed. ns, not significant. **$P < 0.01$.

### Supplementary Fig. S6

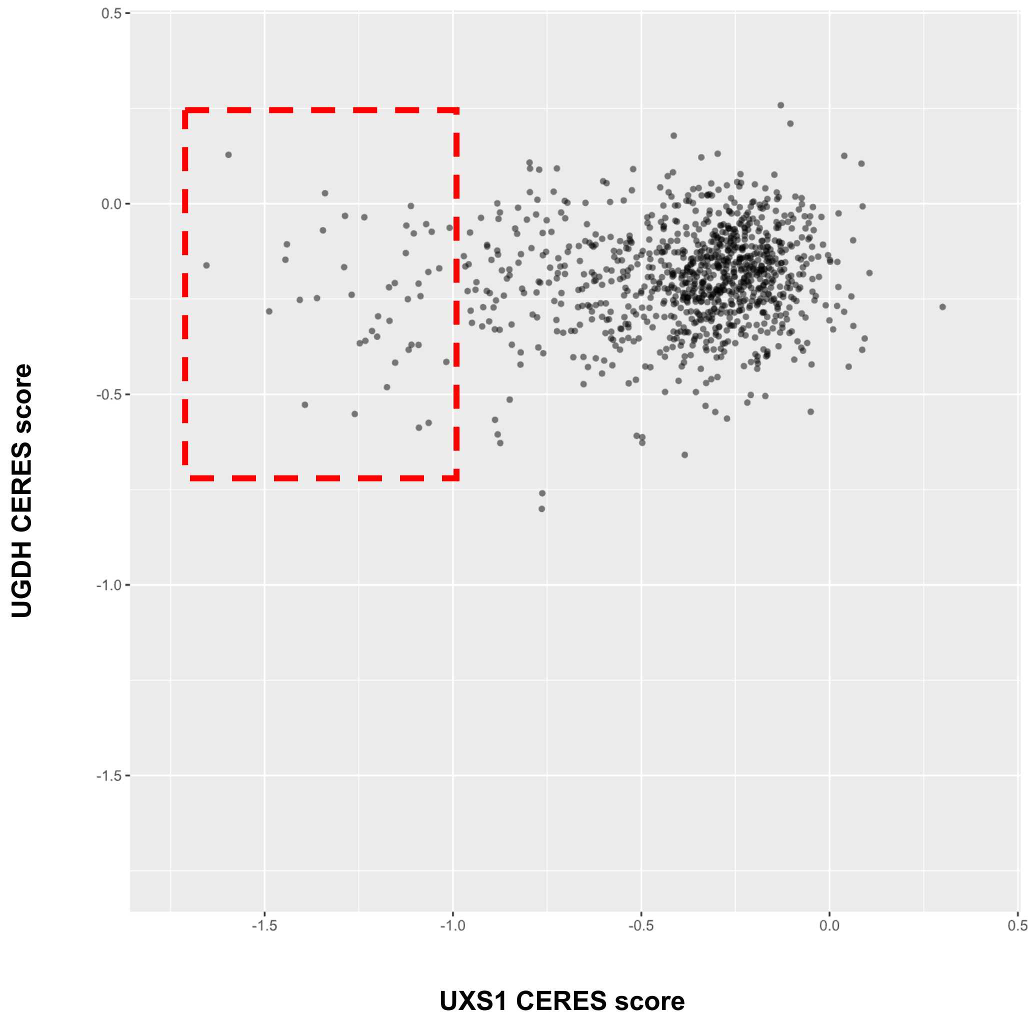


**Supplementary Fig. S6 UXS1-dependent cancer cell lines did not have very negative UGDH CERES scores.** Scatter plot showing the relationship between UGDH CERES score and UXS1 CERES score in cancer cell lines. Cell lines in the red box have UXS1 CERES scores less than -1, which is the median for all common essential genes. Figure was generated with DepMap Public 21Q2 dataset.

**Supplementary Fig. S7**

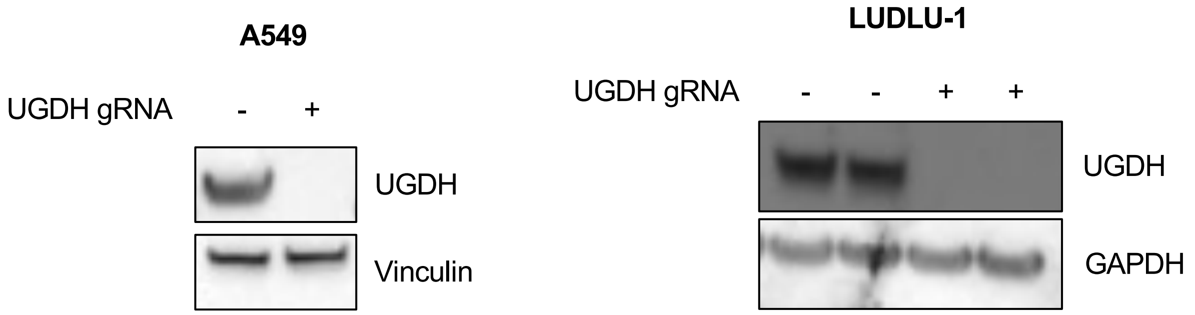


**Supplementary Fig. S7 Validation of UGDH KO cell lines with Western blotting.** Western blotting results of wildtype and UGDH KO A549/LUDLU-1. For LUDLU-1, two replicates of each condition are shown.

**Supplementary Fig. S8**

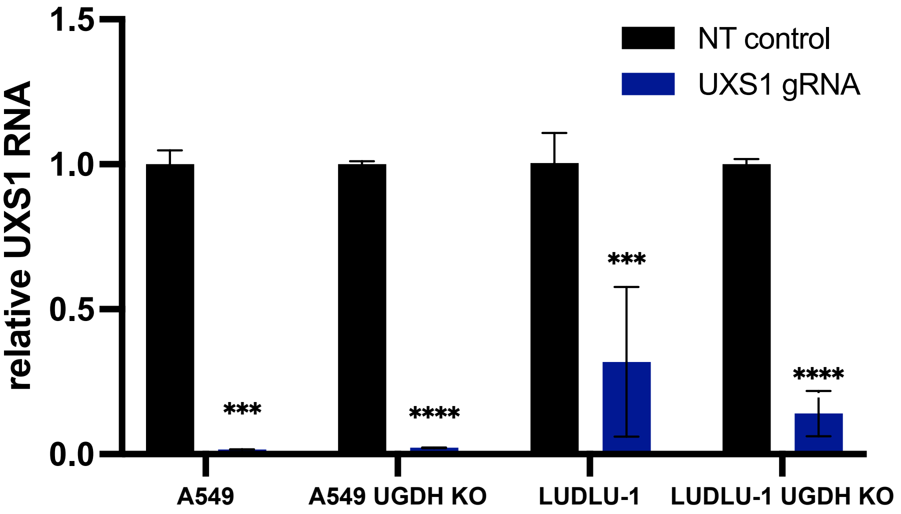


**Supplementary Fig. S8 UXS1 mRNA level was successfully decreased by UXS1 gRNA.** UXS1 mRNA levels were measured by RT-qPCR in A549 or LUDLU-1 cells with/without UGDH stably expressing NT/UXS1 gRNA. Data are mean ± s.d. Two-tailed unpaired Welch's t-test was performed. ***P < 0.001; ****P < 0.0001.

**Supplementary Fig. S9**

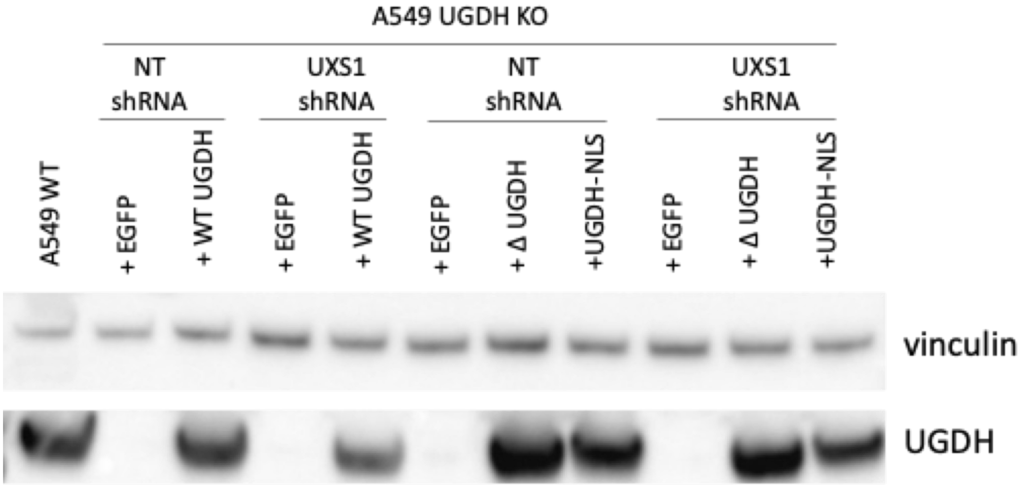


**Supplementary Fig. S9 Validation of exogenous UGDH expression with Western blotting.** Western blotting results of wildtype and UGDH KO A549 cells expressing EGFP/WT UGDH/mutant UGDH/UGDH-NLS and treated with NT/UXS1 shRNA.

### Supplementary Fig. S10

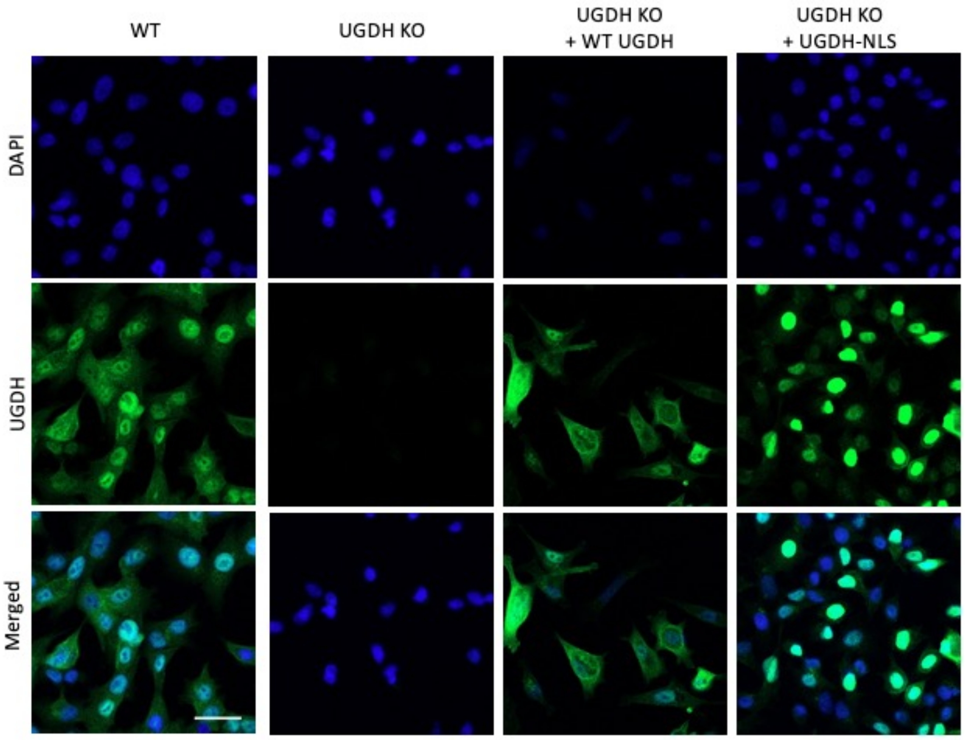


**Supplementary Fig. S10 Subcellular localization of UGDH.** Immunostaining of UGDH in A549/A549 UGDH KO cells expressing WT UGDH/UGDH-NLS. Scale bar: 10 µm.

### Supplementary Fig. S11

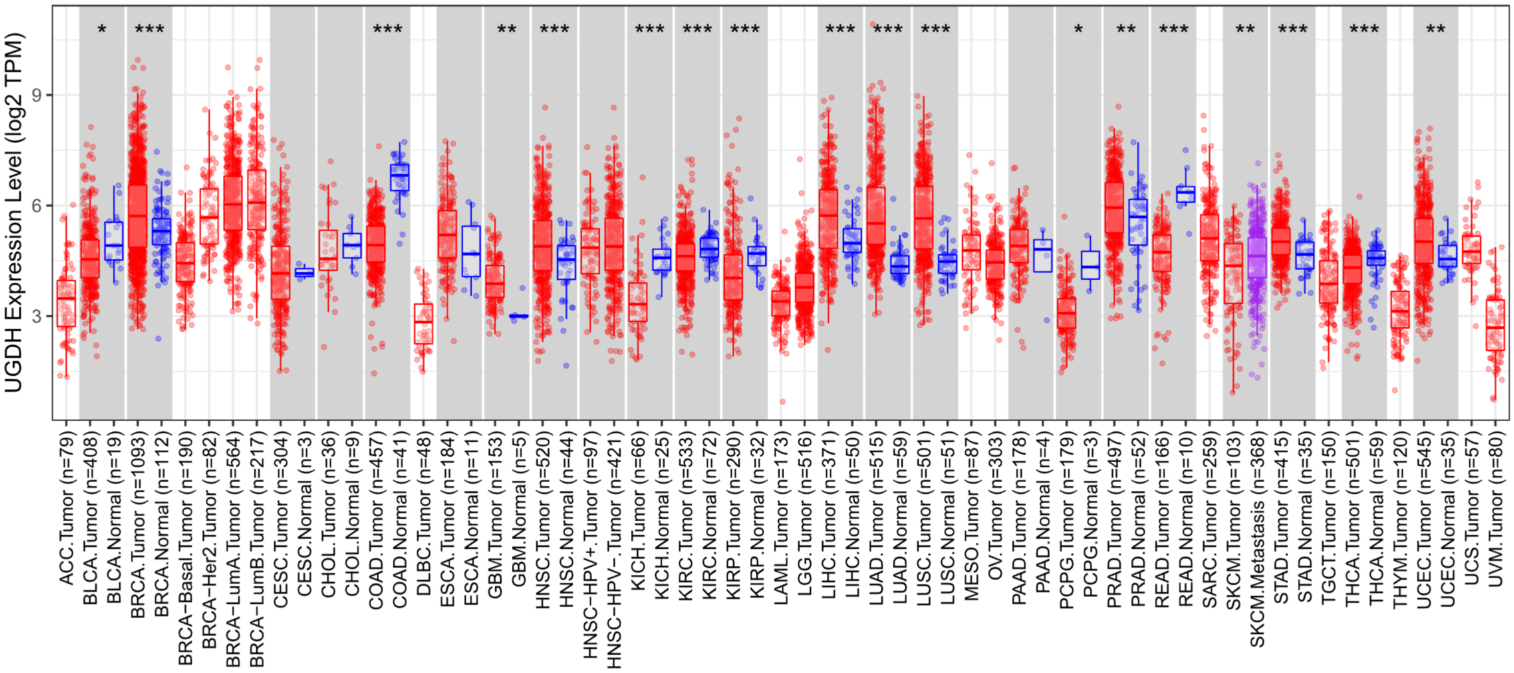

**Supplementary Fig. S11 UGDH expression levels in some cancers are higher than their normal counterparts.** UGDH mRNA levels in different tumor and normal samples. Data are from http://timer.cistrome.org/. Wilcoxon test was performed for statistical analysis (*: p < 0.05; **: p <0.01; ***: p <0.001).

**Supplementary Fig. S12**

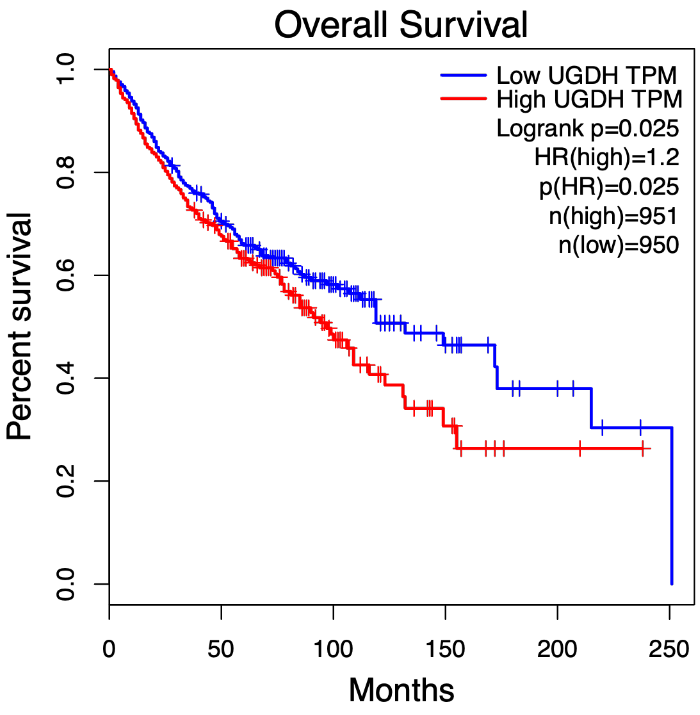


**Supplementary Fig. S12 High UGDH expression is associated with worse overall survival in patients across all cancers, at least during early stages of disease progression.** Data are from http://gepia.cancer-pku.cn/index.html. Overall survival analysis of all types of cancer available in the database. Top 10% of patients with highest UGDH expression levels and bottom 10% were used for the analysis.

### Supplementary Methods

#### RNA purification

Total RNA was isolated from cells with TRIzol reagent following manufacturer's instructions. Briefly, 0.5 M cells were collected. The cell pellet was resuspended in 500 uL TRIzol, and the suspension was incubated for 5 minutes at room temperature. 100 uL chloroform was added and mixed by vortexing. The sample was centrifuged at 12,000 xg at 4 C for 15 minutes followed by RNA precipitation of the aqueous phase with 250 uL isopropanol. The RNA pellet was washed with 1 mL 75% ethanol and resuspended in 30 uL RNase-free water.

#### HPLC-MS/MS analysis

Cells were seeded on 60 mm dish the day before metabolites extraction. On the day, the dish was washed with PBS twice and dipped in liquid nitrogen. Methanol : acetonitrile (50% v:v) was added to the dish and cell extracts were collected with a cell scrapper. Samples were centrifuged at 13,000 rpm at 4 C for 30 minutes. Supernatants were collected and stored at – 80 C until analysis. Standards used were UDP-glucose (Promega, V7091) and UDP-glucuronic acid (Sigma-Aldrich, U6751). The analysis was done on 6495 Triple Quad LC/MS system (Agilent Technologies) with a Ultisil HILIC Amphion, 5um, 2.1×150mm II column (Chromex Scientific, H00274, 31014).

#### Cell lines

HEK293T, NCI-H23, NCI-H1666, NCI-H647 and A549 cells were obtained from ATCC. LUDLU-1 cells were obtained from ECACC. HEK293T and A549 cells were cultured in DMEM supplemented with 10% FBS and 1% penicillin/streptomycin. NCI-H23, NCI-H647 and LUDLU-1 cells were cultured in RPMI supplemented with 10% FBS and 1% penicillin/streptomycin. NCI-H1666 cells were cultured in RPMI supplemented with 5% FBS and 1 % penicillin/streptomycin. All the cells were cultured in a 5% CO2 incubator at 37 C.

#### Antibodies

Antibodies used in this study include UGDH (Atlas Antibodies #HPA036656, 1:2000 for western blot, 1:200 for immunostaining), UGDH (GeneTex #104993, 1:2000 for western blot), vinculin (Invitrogen #700062, 1:1000 for western blot), anti-rabbit secondary (Cell Signaling Technology #7074S, 1:10000 for western blot), anti-rabbit secondary (Thermo-fisher, #A32731) and anti-mouse secondary (Cell Signaling Technology #7076S, 1:10000 for western blot).

#### Gene depletion by CRISPR/Cas9

The gRNA oligos, with sequences for their respective target genes listed below, were annealed and cloned into a lentiCRISPR v2 lentiviral vector. lentiCRISPR v2 was a gift from Feng Zhang (Addgene #52961, #83480). Lentivirus carrying lentiCRISPR v2 plasmid was produced by co-transfecting HEK293T cells with two packaging plasmids, psPAX2 (Addgene #12260) and pMD2.G (Addgene #12259, both were gifts from Didier Trono, and by harvesting viral supernatant after 48 hours by passing through a 0.45 um filter. Collected lentivirus was used directly to infect NCI-H23, NCI-H1666, NCI-H647, LUDLU-1 and A549 cells with the addition of 8 ug/ml polybrene. 48 hours later, the infected cells were selected with puromycin or blasticidin for 5-7 days before being used for subsequent assays.

| | target sequence |
|---|---|
| UXS1 gRNA | GGATTATACATGTAGTTTGG |
| UGDH gRNA | AGACAGAGTACTGATTGGAG |

**Cloning**

UGDH gRNA-resistant transcript sequence was synthesized by Genscript. UXS1 transcript sequence was amplified from cDNA library reversed transcribed from total RNA of NCI-H23. UXS1, WT/enzyme-dead UGDH (C276G) with/without nuclear localization sequence were cloned into pCMV-neomycinR lentiviral vector by using Gibson assembly kit (NEB #E5510S) and site-directed mutagenesis kit (NEB #E0554S). pCMV-neomycinR lentiviral vector was a gift from Huda Zoghbi (Addgene #92194).

**Protein over-expression**

Lentivirus derived from pCMV-neomycinR plasmid expressing tagged UXS1, WT or mutant UGDH and EGFP was produced by co-transfecting HEK293T cells with two packaging plasmids (psPAX2 and pMD2.G) and by harvesting viral supernatant after 48 hrs by passing through a 0.45 um filter. Collected lentivirus was used directly to infect A549 cells with the addition of 8 ug/ml polybrene. 48 hrs later, the infected cells were selected with 20 uL/mL G418 solution (Roche, 4727894001 ) for 5-7 days before being used for subsequent assays.

**Gene knockdown by shRNA**

UXS1 shRNA oligos (GCTTGCTTTAATGAAATGGAT) was annealed and cloned into a pLKO.1-Puromycin lentiviral vector (Addgene #10878, a gift from David Root). Lentivirus carrying pLKO.1 plasmid was produced by co-transfecting HEK293T cells with two packaging plasmids (psPAX2 and pMD2.G) and by harvesting viral supernatant after 48 hrs by passing through a 0.45 um filter. Collected lentivirus was used directly to infect NCI-H23 and A549 cells with the addition of 8 ug/ml polybrene. 48 hrs later, the infected cells were selected with 1 ug/mL puromycin for 5-7 days before being used for subsequent assays.

**Cell growth assay (CyQUANT)**

3500 NCI-H23/NCI-H1666/NCI-H647/LUDLU-1/A549 cells were seeded in each well of 96-well plate for cell growth assay in 200 uL volume. 8 days later, relative cell number was assessed by using the CyQUANT NF cell proliferation kit (Invitrogen, C35006) according to the manufacturer's instructions.

**Cell growth assay (BrdU)**

BrdU cell growth assay was performed with BrdU assay kit (Merck, #QIA58) according to the manufacturer's instruction. In brief, 1 x 10^4 cells NCI-H23/A549 cells were seeded in each well of 96-well plate for cell growth assay in 100 uL volume followed by adding 20 uL BrdU label. 24 hours later, the plate was fixed and stained with BrdU antibody.

**RT-qPCR**

The extracted RNA was reversely transcribed into cDNA using the PrimeScript™ RT Master Mix (Perfect Real Time) (Takara #RR036A) according to the manufacturer's instructions. The obtained cDNA samples were diluted and used for real-time quantitative PCR (RT-qPCR). Power SYBR green PCR master mix (Applied Biosystems #4367659) and gene specific primers with sequences listed below were used for PCR amplification and detection on a QuantStudio3 real-time PCR system (Applied Biosystems). The RT-qPCR data were normalized to GAPDH or b-actin and presented as fold changes of gene expression in the test sample compared to the control.

| | sequence |
|---|---|
| UXS1_Forward | GCTCGTGACCATTTCCAAGG |
| UXS1_Reverse | TTCGGAATTTCACCAGAAGGA |
| UGDH_Forward | AGCCTCCCCTCCAAACTACA |
| UGDH_Reverse | GGCCACTCGCACTTCCAC |

**Immunostaining**

Cells were seeded on cover-slips coated with 0.01% poly-L-lysine the day before staining. On the day, cells were fixed with 2% Paraformaldehyde and permeabilized with 0.2% Triton-X. Samples were then washed and stained with UGDH antibody overnight followed by secondary anti-rabbit 488A antibody. Slides were visualized with LSM980 confocal microscope (Zeiss). Images were analyzed with ImageJ.
